# The replicative amplification of MITEs and their impact on rice trait variability

**DOI:** 10.1101/2020.10.01.322784

**Authors:** Raul Castanera, Pol Vendrell-Mir, Amélie Bardil, Marie-Christine Carpentier, Olivier Panaud, Josep M. Casacuberta

## Abstract

Transposable elements (TEs) are a rich source of genetic variability. Among TEs, Miniature Inverted-repeat Transposable Elements (MITEs) are of particular interest as they are present in high copy numbers in plant genomes and are closely associated with genes. MITEs are deletion derivatives of class II transposons, and can be mobilized by the transposases encoded by the latters through a typical cut-and-paste mechanism. However, this mechanism cannot account for the high copy number MITEs attain in plant genomes, and the mechanism by which MITEs amplify remains elusive.

We present here an analysis of 103,109 Transposon Insertion Polymorphisms (TIPs) in 1,059 *O. sativa* genomes representing the main rice population groups. We show that an important fraction of MITE insertions has been fixed in rice concomitantly with rice domestication. However, another fraction of MITE insertions is present at low frequencies. We performed MITE TIP-GWAS to study the impact of these elements on agronomically important traits and found that these elements uncover more trait associations than SNPs on important phenotypes such as grain width. Finally, using SNP-GWAS and TIP-GWAS we provide evidences of the replicative amplification of MITEs, suggesting a mechanism of amplification uncoupled from the typical cut-and-paste mechanism of class II transposons.

## Introduction

Transposable Elements (TEs) are an essential component of plant genomes. Their capacity to amplify and create new genetic variability by insertion/excision, and the possibility that their multiple copies offer for recombination, make them a rich source of genomic variants that can be selected through evolution (Tenaillon *et al*., 2010). TE-induced mutations include gene knock-outs, but also the induction of gene epigenetic silencing or changes in gene regulation by inactivating enhancers or repressors upon insertion or by adding new regulatory elements contributed by the TE (Lisch, 2013). Miniature Inverted-repeat Transposable Elements (MITEs) are short non-coding TEs thought to be deletion derivatives of class II cut-and-paste TEs (Feschotte *et al*., 2002). However, differently to the class II TEs they derive from, MITEs are found in high-copy number in plant genomes (Chen *et al*., 2014). The apparent contradiction between a conservative cut-and-paste mechanism of transposition and MITEs high copy number was highlighted short after MITE discovery (Feschotte *et al*., 2002; Casacuberta and Santiago, 2003), but the mechanism by which MITEs amplify is still obscure. MITEs are tightly associated with plant genes (Santiago *et al*., 2002; Lu *et al*., 2012; Benjak *et al*., 2009), and examples of these elements potentially altering gene expression have accumulated over the years (Santiago *et al*., 2002; Lu *et al*., 2012; Naito *et al*., 2009; Yang *et al*., 2005; Xu *et al*., 2020; Zheng *et al*., 2019; Yin *et al*., 2020). Moreover, it has recently been shown that MITEs frequently contain Transcription Factor Binding Sites (TFBS) in plants, which may allow their mobilization to alter transcriptional networks by rewiring new genes (Morata *et al*., 2018), and that they can induce structural variability through aberrant transposition events (Chen *et al*., 2020).

In the last few years, different methods for analyzing TE insertion polymorphisms (TIPs) in resequenced genomes have been developed (Vendrell-Mir *et al*., 2019), and LTR-retrotransposon (LTR-RT) TIPs have recently been used to perform GWAS to study LTR-RT dynamics in rice (Carpentier *et al*., 2019), and the genetic basis of agronomic traits in rice and tomato (Akakpo *et al*., 2020; Domínguez *et al*., 2020). These recent publications show that LTR-RT TIPs can allow discovering associations not seen with conventional GWAS strategies based on SNPs and highlight the importance of analyzing the fraction of the genetic variability TEs account for. MITEs have not been used in TIP-GWAS as they were considered as more difficult to analyze due to their small size and high copy number (Domínguez *et al*., 2020). However, recent benchmarking efforts of TIP prediction tools have allowed us to propose efficient approaches to analyze MITE insertions (Vendrell-Mir *et al*., 2019). Here we used these approaches to define MITE TIPs in 1,059 rice varieties and used them to perform TIP-GWAS on different agronomic traits. We show that, indeed, MITE TIPs reveal an additional fraction of the genetic variability associated to important crop traits and may allow discovering the underlying causal genes for some traits. In addition, we used MITE TIP-GWAS to study MITE dynamics in rice highlighting the impaact of MITEs on rice domestication and breeding and confirming that MITEs amplify by a replicative mechanism from a reduced number of MITE copies.

## Results

### 1. TE annotation and selection of rice varieties

We performed a stringent TE annotation using the REPET package (Flutre et al, 2011). As MITEs are particularly prevalent in plants (Chen *et al*., 2014), we complemented this annotation with a targeted annotation of MITEs based on MITE-hunter (Han et al, 2010) and the already published annotations in the P-MITE database (Chen et al, 2014). We used this pipeline to annotate 3 assembled rice reference genomes belonging to the three major rice subgroups (Indica, Japonica and Aus), and we clustered all TE consensuses to obtain a global TE library of the species. We identified a total of 821 complete TE families belonging to all major TE orders, excluding incomplete and chimeric elements (Supporting Figure 1, Supporting dataset S1). In spite of the stringency of the approach used to build the TE library, a RepeatMasker (http://www.repeatmasker.org) annotation with the 821 consensus sequences on the Nipponbare reference genome (MSU7) (Kawahara *et al*., 2013) showed highly congruent results in comparison to recently published rice TE annotations. More specifically, 95% of the annotation overlaps rice 6.9.5.liban annotation (Ou *et al*., 2019) (78% overlap in the opposite direction), and the classification agreement at the order level between reciprocal consensuses of the two libraries was 99%. However, the annotation used here has fewer LTR-RT consensuses (112 vs 389), probably due to the high stringent approach that did not retained partial, low-copy number and degenerated LTR-RT families, and has much more MITE consensuses (400 vs 173) due to the dedicated tools used.

In order to study the TE dynamics in rice, and in particular that of MITEs, we took advantage of the availability of resequencing data of 3000 rice varieties to look for TE insertion polymorphisms (TIPs) in the different varieties of the TEs annotated in the reference genomes. A recent benchmarking exercise indicated that PopoolationTE2 (Kofler *et al*., 2016) is a good tool for this purpose, but also that coverage is a key factor than can limit the detection of TIPs (Vendrell-Mir *et al*., 2019). For this reason, we selected the 1,059 rice genomes sequenced at 15x or more (Supporting Table S1) and subsampled all genomes to 15x in order to be able to perform an unbiased comparisons of TIP abundance between varieties.

We found an average of 40,568 insertions per variety. We filtered the insertions using a zygosity cut-off of 0.7 as recently recommended to avoid false positive calls (Vendrell-Mir *et al*., 2019). After applying this filter, we obtained an average of 18,463 insertions per variety.

The number of TIPs per variety is variable (Figure 1), but a first analysis indicated that CAAS varieties had a significantly lower number of TIPs than IRIS varieties (Supporting Figure S2), despite the identical processing pipeline followed to sequence the selected varieties (Li *et al*., 2014). Although we could not identify the reason for such a difference, we warn that it could be a limitation for quantitative studies targeting TIPs in the 3,000 genomes dataset. In order to avoid any possible bias, we decided to use only the IRIS data, which reduced the number of rice varieties analyzed to 738.

**Figure 1.**
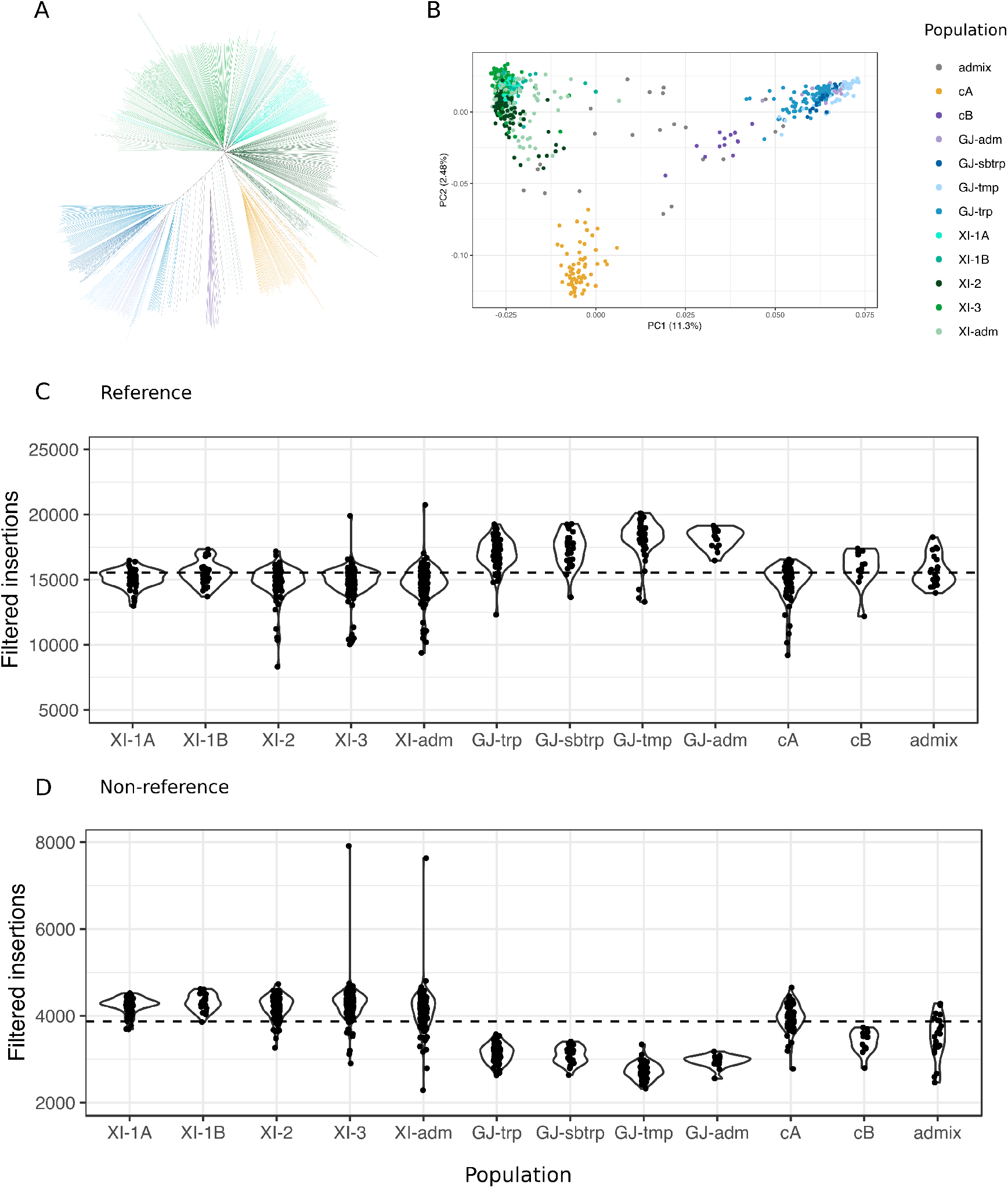
TIP-based population structure and TIP content in 738 rice varieties. A) Neighbor-joining tree. B) PCA based on a presence/absence matrix of 103,109 insertion polymorphisms. Colors denote the nine SNP-based sub-populations described by Wang et.al (2018). Reference (C) and non-reference (D) insertions per variety grouped by subpopulation.

### 2. TE dynamics in rice varieties

We detected a total of 103,109 TIPs in the 738 rice varieties corresponding to 32,449 retrotransposons and 70,660 DNA transposons of all orders. PCA analysis and phylogenetic trees performed based on predicted TIPs are congruent with the previously defined groups of rice varieties based on SNPs (Wang *et al*., 2018) (Figure 1A, Figure 1B). Although the analyses using only the TIPs of each type of TEs gave similar results, the resolution of the different groups varied among them (Supporting Figure S3). For example, the analysis based on LTR-RT TIPs better resolved all the different rice groups as compared with MITEs, probably due to a higher number of non-fixed and non-private LTR-RT TIPs.

The analysis of TIPs shows that most TE families have a relatively constant number of TIPs per variety among the four main varietal groups (Japonica, Indica, Aus and Aro), with a Coefficient of Variation between groups (CV) lower than 20% in 82% of the families, and with a median of 9.5% (Figure 2A). However, a few number of families are much more variable, the CV reaching a maximum of 128.2%. Up to 150 TE families had a coefficient of variation higher than 20% and have potentially experienced a varietal group-specific amplification (Figure 2A, 2B). The CV as well as the proportion of private insertions, are higher for smaller families (Figure 2C, 2D), suggesting that these families have been more active recently. Among the different TE orders, LTR-retrotransposons are the only order significantly enriched among the group-specific amplification families (Fisher’s exact test p-value = 0.0062, Table 1).

**Figure 2.**
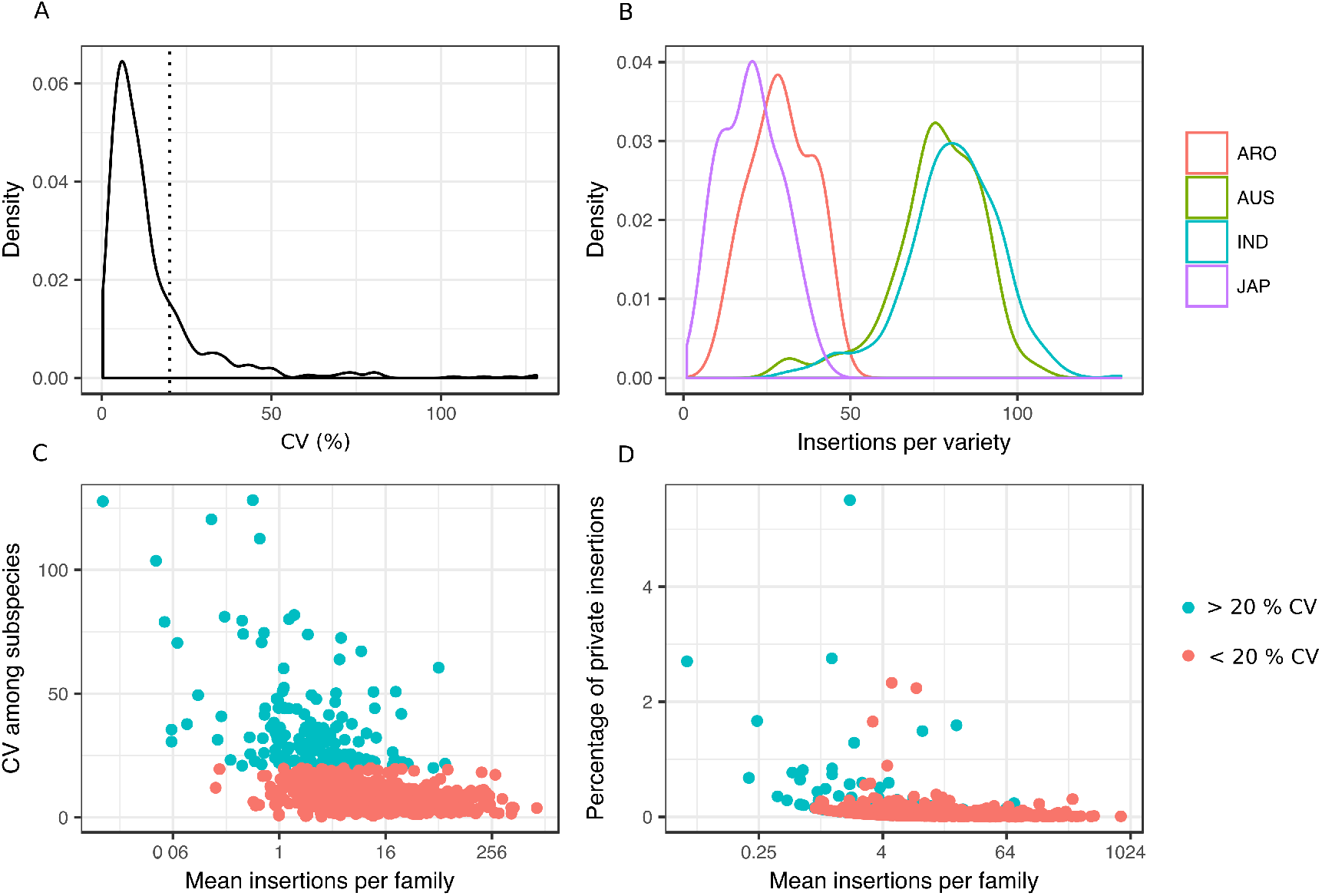
Signatures of subspecies-specific activity in rice TE families. A) Distribution of inter-subspecies Coefficient of Variation in the average family insertion number. Dotted line represents a CV of 20%. B) Example of a gypsy LTR-retrotransposon family (RLX_comp_MH63_B_R175_Map20) showing signs of amplification in Indica and Aus subspecies after their split with Japonica and Aro. C) Mean insertion per family in population *versus* CV. D) Mean insertion per family in population *versus* the number of private insertions.

**Table 1.** Number of TE-families per TE order showing group-specific amplification. The number of families showing a coefficient of variation among varietal groups (CV) of more than 20% are shown with respect to the total number of families for each specific TE order. Only the LTR-RT order is significantly enriched (Fischer’s exact test) in families that show group-specific amplification (shown in bold)

| Order | CV > 20 % | N. families | % | pvalue |
| --- | --- | --- | --- | --- |
| Helitrons | 1 | 29 | 3.45 | 0.0708 |
| TIR-TEs | 34 | 168 | 20.24 | 0.5843 |
| MITEs | 62 | 400 | 15.50 | 0.1094 |
| LINEs | 19 | 104 | 18.27 | 1,00 |
| LTR-RTs | 34 | 112 | 30.36 | **0.0062** |
| SINEs | 0 | 8 | 0.00 | 0.6171 |

In order to get more insight on the dynamics of the different types of TEs we analyzed the TIP frequency distribution for each type of TEs in a subset of 382 varieties that correspond to traditional varieties and are a good representation of the variability of the species (Gutaker *et al*., 2020). As shown in Figure 3A, the frequency distribution is different for different orders of TEs. In particular, and as previously shown (Carpentier *et al*., 2019), our study reveals that most LTR-RT TIPs are present at very low frequency (LF), which could suggest both a recent activity and a high turnover of LTR-RT insertions. On the contrary, although there is an important number of MITE TIPs present at low frequency, there is a significant number of them that appear to be fixed in the population, presenting a “U-shape” frequency spectrum with an excess of high-frequency (HF) derived mutations at the expense of middle-frequency variants. This suggests that, whereas most LTR-RT insertions are recent, rice has retained an important fraction of older MITE insertions. This can also be seen when analyzing the fraction of TIPs corresponding to insertions present in rice (*Oryza sativa* ssp. *Japonica* cv. Nipponbare) and nine different *Oryza* genomes (Zhou *et al*., 2020) (Figure 3B). Whereas less than 10% of the LTR-RT insertions present in rice (Nipponbare) are also present in *Oryza glumaepatula*, more than 30% of the MITE insertions are also present in this relatively distant *Oryza* genome (Figure 3B). This data clearly suggests that an important fraction of MITE insertions have been fixed during rice evolution. As MITEs tend to concentrate close to genes (Casacuberta and Santiago, 2003; Feschotte *et al*., 2002) these fixed insertions may have played an important role in the evolution of rice genes. On the other hand, an analysis of the frequency within rice varieties of the insertions shared between Nipponbare rice and *O. glumaepatula*, shows that these relatively old insertions have been maintained in the whole population (Figure 3B), confirming that, as already proposed (Feschotte *et al*., 2002), most MITE insertions are highly stable, in spite of the possibility of being excised by related transposases (Chen *et al*., 2020).

**Figure 3.**
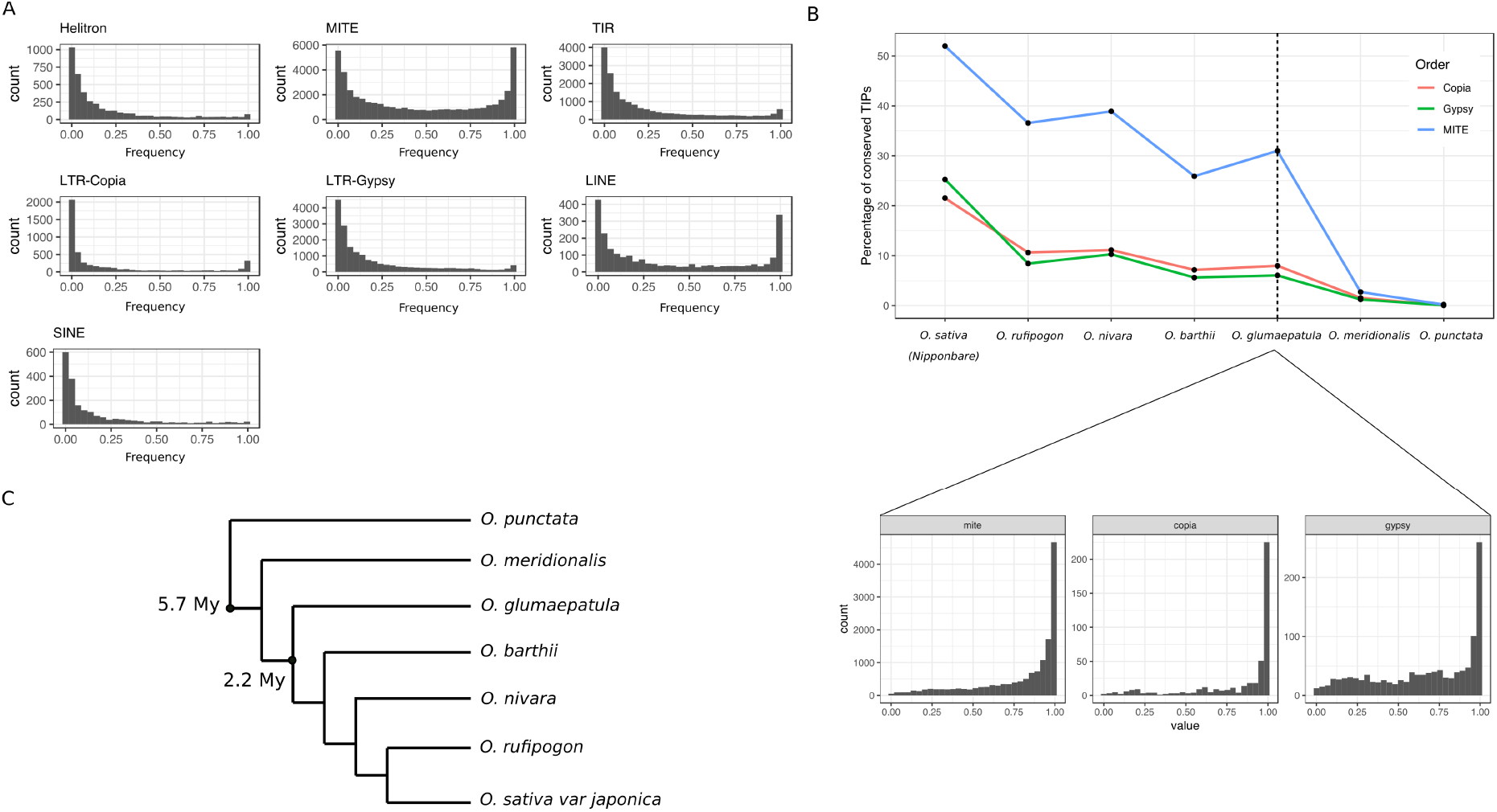
TIP population frequencies and their conservation in wild *Oryza* species. A) TIP frequencies in 382 rice traditional varieties. Number of TIPs per order are: 4,113 Helitrons, 45,837 MITEs, 18,592 TIR-TEs, 5,026 copia LTR-RTs, 18,742 gypsy LTR-RTs, 2,402 LINEs and 2,000 SINEs. B) Percentage of Population TIPs present in *O. sativa* ssp. *Japonica* cv. Nipponbare and in 6 wild *Oryza* species. Lower panels show the frequency of the TIPs conserved between *O. sativa* ssp. *Japonica* cv. Nipponbare and *O. glumaepatula* in the traditional rice varieties. C) Dendrogram representation of the phylogenetic relationships between the *Oryza* species analyzed. Branch lengths do not represent real phylogenetic distances. Divergence times were obtained from http://www.timetree.org/.

An analysis of the chromosomal distribution of MITE TIPs shows that it is similar to that of copia LTR-RTs and also follows that of genes (Figure 4A and Supporting Figure S4). This is expected as copia LTR-RTs and MITEs are known to be closely associated to plant genes (Casacuberta and Santiago, 2003). However, a comparison of the distribution of TIPs present at low- and high-frequency (1^st^ and 4^th^ frequency quartiles, respectively), which allow starting to discriminate between the role of target specificity and selection in shaping TE distribution, shows clear differences between the two TE groups. Whereas only low frequency copia LTR-RTs insertions follow gene distribution, MITE insertion association with genes is stronger for high frequency insertions (Figure 4, Supporting Figure S5). This suggests that whereas in general both copia LTR-RTs and MITEs seem to target genic regions for integration, LTR-RTs are progressively cleaned away from these regions whereas MITEs inserted close to genes are more frequently maintained. A detailed analysis of the non-reference insertions with respect to genes shows that MITEs concentrate in the 5’ and 3’ proximal regions (<500 nt) of genes (Figure 4C, Supporting Figure S6A). MITE enrichment in the gene space was stronger in the 5’ upstream regions and was found for both high frequency and low frequency insertions (Supporting Figure S6B). Therefore, MITEs could have had an important impact on gene variability in the recent evolution of rice. As shown in Figure 3B, although a significant fraction of MITE insertions are present in the rice wild relative *Oryza rufipogon*, more than 60% of them seem specific of domesticated rice. We analyzed the MITE insertions that are present at HF in rice varieties and are absent from *O. rufipogon* as they could potentially be linked to domestication mutations, and found that the fraction of insertions close to genes is significantly higher than for the whole MITE insertions here described (63% vs 54%). Among the 834 genes that present a MITE insertion present at HF in domesticated rice and absent from wild rice, there are some already characterized rice genes with important functions during development or under stress conditions and that may be related to important agronomic traits (Supporting Table S2).

**Figure 4.**
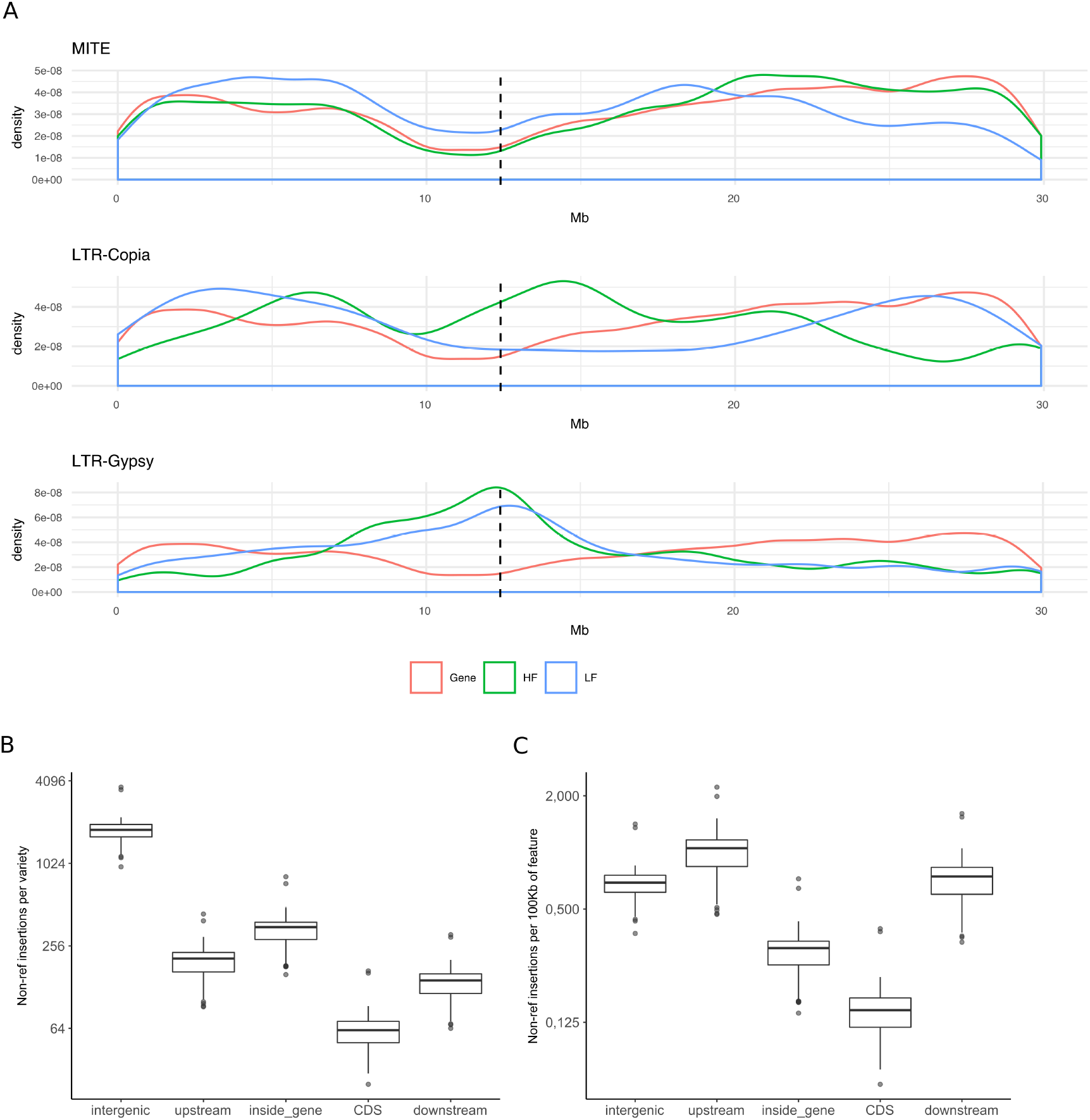
Genome-wide distribution of TIPs and their impact on genes. A) Density plot showing the chromosomal distribution of High-frequency (HF) and Low-Frequency (LF) TIPs of MITEs, gypsy and copia LTR-RTs and genes (Chr04, full genome shown in Supporting Figure S5). B) Number of non-reference MITE insertions per variety. C) Number of non-reference MITE insertions per 100Kb of genomic features.

### 3. Association of MITE insertions with trait variability

Genome-wide association studies (GWAS) using LTR-RT TIPs as a genotype, instead of SNPs, have recently been performed in rice and tomato (Domínguez *et al*., 2020; Akakpo *et al*., 2020). These studies showed that these TIPs can reveal additional genetic associations with traits that are not seen with SNPs. Here we wanted to explore the potential of MITE TIPs for TIP-GWAS. We reasoned that their small size, high copy number, and close association with genes, could make MITEs particularly suited for this purpose. We performed GWAS using MITE TIPs as genotype and the phenotypic data available for the 451 Indica varieties (Mansueto *et al*., 2017; Jackson, 1997). We obtained significant associations between specific MITE insertions and six phenotypes (grain width, length and weight, salt injury at EC18, flowering time and leaf length, Supporting Tables S3 and S4). Some of the peaks obtained where coincident with peaks obtained using SNPs as genotype, suggesting that both were in Linkage Disequilibrium (LD) with the causal mutation (that could also be the SNP or the MITE TIP themselves). However, in many cases the MITE TIP-GWAS revealed different peaks allowing to explore additional genetic regions potentially linked to trait variability (Supporting Tables S3 and S4).

As an example, whereas the SNP-GWAS revealed a single genomic locus linked to grain width, the MITE TIP-GWAS revealed nine different significant association peaks (Figure 5, Supporting Table S3). The only SNP association obtained was found at the well-characterized GSE5 gene of the GW5 locus controlling grain size (Duan *et al*., 2017). The leading SNP was located at 3.9 Kb of the GSE5 gene. Interestingly, the strongest association using MITE TIPs, MITE-1 (p-value = 3.28e^-25^) (Figure 5) corresponded to a MITE insertion located 350 nt upstream of the GSE5 gene. This insertion is in strong LD with a deletion located 4,500 bp upstream the GSE5 gene, and the co-occurrence of these two structural variants constitute one of the three main haplotypes of GSE5 in cultivated rice (GSE5^DEL1+IN1^) (Duan *et al*., 2017). However, in spite of the MITE insertion being closer to the GS5 gene than the deletion, our analysis suggest that the MITE insertion is not the causal mutation responsible of grain width, as the few varieties carrying the MITE but not the deletion have narrow grains (Supporting Figure S7).

**Figure 5.**
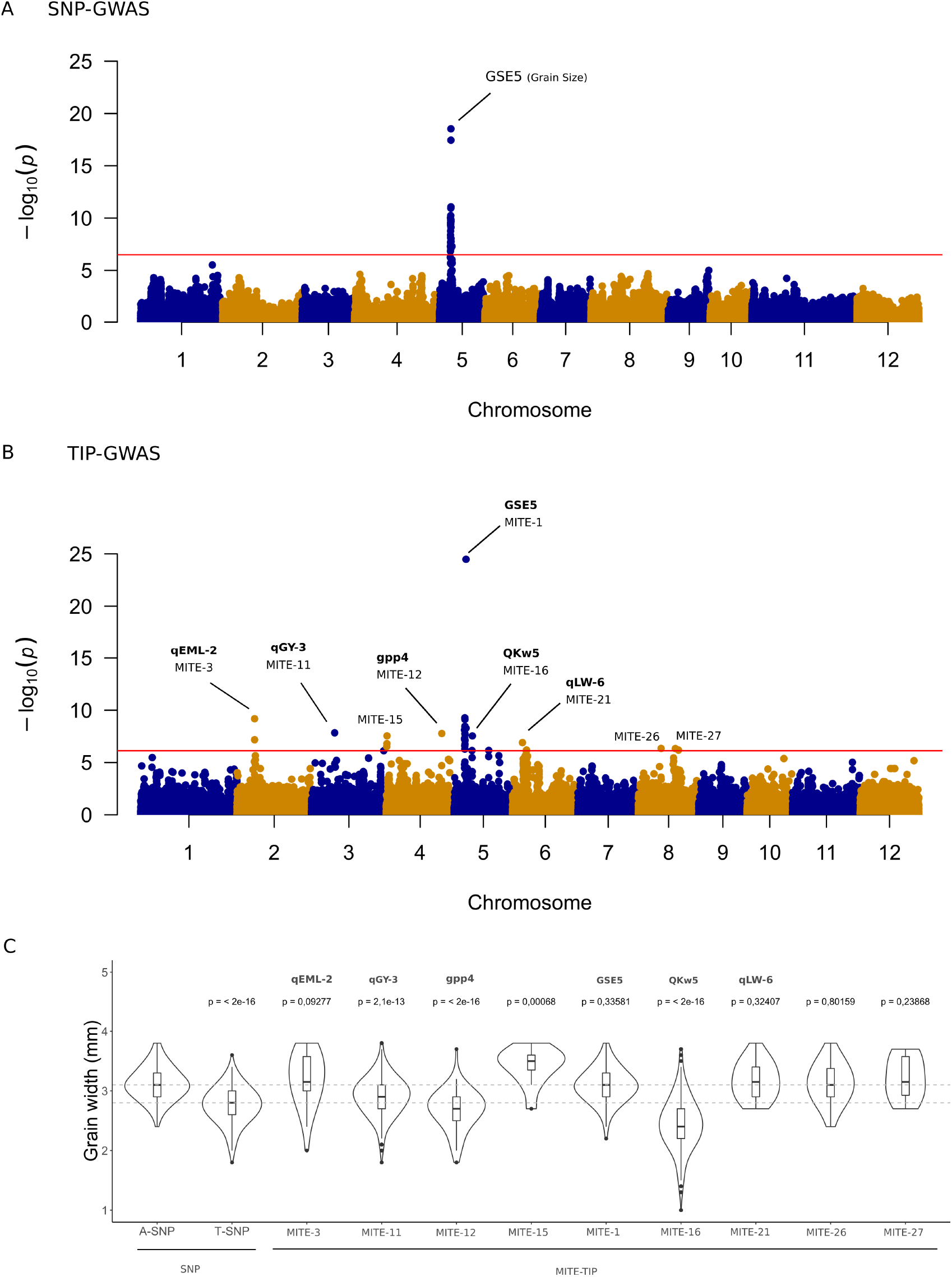
Association of MITE transposon insertions with higher rice grain width. A) Rice grain width SNP-GWAS using 451 rice Indica varieties. B) Rice grain width TIP GWAS using MITEs and TIR-TE insertion polymorphisms as genotype (66,368 TIPs in total). Red line represents the Bonferroni-adjusted significant threshold (3.2e^-07^). Significant MITE TIPs overlapping known seed QTLs are marked in the manhattan plots. C) Violin plots showing the rice grain width in the different subsets of varieties carrying each variant of the leading SNP at position chr05:5361195 (left panel), coinciding with the upstream region of the seed gene GSE5. Rice grain width of varieties carrying every significant TIP are shown in the right panel. Differences between each distribution and the control (Genotype “A” at chr05: 5361195) were tested using a two-tailed Wilcoxon rank sum test (p-value cutoff = 0.05). More information about the significant MITE-TIPs is shown in Supporting Table S3.

In addition to the association related to the GSE5 gene, we detected eight more peaks with MITE insertions associated with grain width, several of which are located close to genes or within QTLs already characterized as linked to grain phenotypes. The MITE-15 insertion is located at 33 Kb of the OsARG gene, which controls grain yield in rice (Ma *et al*., 2013), and five additional MITE TIPs are within a previously characterized QTLs linked to grain yield (MITE-11 at qGY-3 (Mao *et al*., 2003) and MITE-21 at QTARO QTL-450 (Yonemaru *et al*., 2010)), embryo length (MITE-3 at qEML-2 (Dong *et al*., 2003)), grains per panicle (MITE-12 at gpp4 (Xiao *et al*., 1996)), grain shape (MITE-21 at qLW-6 (Yan *et al*., 2003)) and 1000 kernel weight (MITE-16 at QKw5 (Li *et al*., 1997)), which reinforces de relevance of the associations found. A detailed analysis of the MITE insertion positions showed that four out of these nine significant MITE TIPs were located inside genes or in their close vicinity (1000bp upstream or downstream) (Supporting Table S3). We anticipate that these genes could be good candidates to be linked to the grain width trait.

### 4. Genetic factors linked to MITE amplification

In addition to study the genetic basis of agronomic traits, GWAS can also be used to study the genetic determinants of TE activity, as recently done for LTR-RTs in rice (Carpentier *et al*., 2019). Here we use this approach to study the genetic determinants of MITE amplification. This is particularly relevant as although MITEs are known to be mobilized by transposases of class II related elements (TIR-TEs), a canonical cut-and-paste mechanisms does not allow for the high-copy numbers that MITEs frequently attain in eukaryotes, and particularly in plant genomes (Chen *et al*., 2014). Nevertheless, in searching for genetic determinants of MITE amplification, we paid particular attention to the possible role of transposases related to the MITE families analyzed.

We performed a GWAS analysis on 451 rice varieties belonging to Indica subspecies using a SNP matrix of 173,260 SNPs as genotype, and the MITE copy number of 400 MITE families as phenotype, running one association analysis per TE family. We used Indica populations as they are present in a higher number in our dataset, which provide us with more power for the GWAS analyses. As a first approach, and in order to directly check for the potential role of transposase-encoding TEs related to MITE families in their amplification, we analyzed 18 MITE families showing significant sequence similarity with TIR-TEs from which they probably derive (Supporting Table S5). We identified peaks of SNPs significantly associated with the copy number of seven of those 18 MITEs families. None of these SNPs peaks had a TIR-TE insertion with significant sequence similarity with the MITE family at less than 100kb. Interestingly, five of the associated SNPs peaks (for four families) had an insertion of a MITE of the same family at less than 100kb (Supporting Table S5). As a complementary approach, we performed TIP-GWAS using TIR-TE and MITE TIPs as genotype. TIR-TE TIP-GWAS did not reveal any association of a particular TIR-TE with the copy number of the related MITE family. On the contrary, a TIP-GWAS performed with MITE TIPs revealed significant associations of a MITE insertion with the MITE family copy number for 11 out of 18 MITE families, including the four families for which we detected an associated SNP peaks with a MITE of the same family located close. As an example, Figure 6 shows the analysis of the genetic factors associated with the copy number of the MITE family SE260300235fam318_632. The SNP-GWAS revealed two different SNP peaks strongly associated with the MITE family copy number (Figure 6A). MITE TIP-GWAS revealed several peaks of MITE TIPs significantly associated with the MITE family copy number (Figure 6B), and in all cases the leading TIP corresponded to a MITE of the same family (in green in Figure 6B). The leading TIPs of chromosomes 2 and 6 are in strong LD with the corresponding leading SNPs of the associations detected by SNP-GWAS (Figure 6E), and their presence strongly correlates with an increase of copy number of the MITE family (Figure 6D). On the contrary, the TIR-TE TIP-GWAS did not reveal any association of the TIR-TE family closely related to the MITE family (see Figure 6C), the hAT family DTX_comp_IRGSP_B_R1932, with the amplification of the MITE family (Figure 6F).

**Figure 6.**
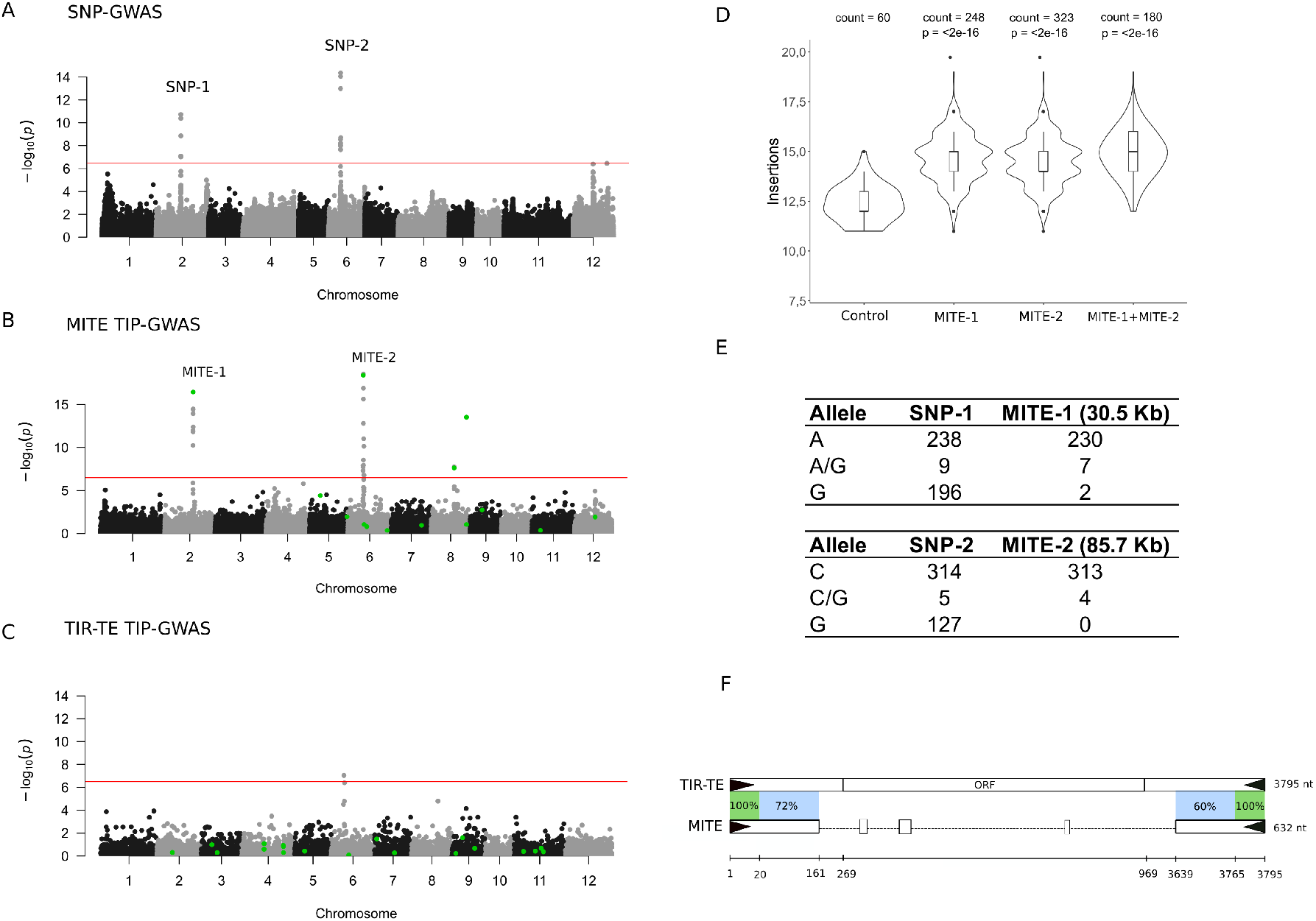
Identification of MITE copies as main genetic factors responsible for their amplification in Indica rice. A) SNP-GWAS analysis of MITE family SE260300235fam318_632 using insertion numbers as phenotype and SNPs as genotype (451 Indica varieties). B) TIP-GWAS analysis using insertion numbers as phenotype and all MITE TIPs as genotype. C) TIR-TE TIP-GWAS analysis using insertion numbers as phenotype and all TIR-TE TIPs as genotype. In green, TIPs from the family SE260300235fam318_632 (panel B) and from the distantly related TIR-TE DTX_comp_IRGSP_B_R1932 (panel C). Red line represents the Bonferroni-adjusted significant threshold (3.2E-07). D) Mean MITE insertion numbers of varieties containing or not the associated MITEs. Control are all the varieties that do not contain any of the two MITEs. The number of varieties present in each category are indicated upstream each violin. P-values correspond to two-tailed Wilcoxon rank sum tests comparing each category *vs* the control category. E) Genotypes of the leading SNPs in all Indica varieties and in the subsets of varieties carrying the two associated MITEs. Distances between each MITE and the leading SNP is reported in Kb. F) Schematic representation of the nucleotide conservation between the consensus sequences of the full TIR-TE element DTX_comp_IRGSP_B_R1932 and its related MITE SE260300235fam318_632. ORF = Open Reading Frame.

We extended this analysis to the 400 MITE families identified here. We found significant association SNP peaks for 175 out of the 400 MITE families. The SNPs peaks associated with MITE family amplification were distributed along the 12 chromosomes without a particular enrichment in any genomic region (Supporting Table S6. None of these associated SNPs had a TIP related to a TIR-TE with similarity (even limited similarity at the TIR level) to the corresponding MITE family. On the contrary, up to 37% of these peaks (66) have a MITE TIP of the same family at less than 100kb (Supporting Table S7). In 96% of the cases, the presence of this MITE copy is linked to an increase of MITE copy number of the respective. We used three assembled Indica varieties from the recently published platinum genomes (Zhou *et al*., 2020) to analyze the insertions corresponding to the TIPs located close to the 66 SNP peaks associated to MITE copy number and in 48 out of 49 cases where the TIP is predicted to be present in one of the platinum genomes, we confirmed the presence of the MITE insertion in the genome assembly..

As a complementary approach, we performed GWAS using MITE TIPs or TIR-TE TIPs instead of SNPs as genotype. We detected associations of a MITE TIP with the copy number of 301 different MITE families (75% of them). In most of these cases (186) the most significant MITE TIP belongs to the same MITE family for which they are potentially affecting the copy number. These associations include 44 of the 66 associations found previously by SNP-GWAS. The high number of associations with MITE insertions of the same family strongly suggests that the association of the MITE is not the result of a LD with another genetic factor influencing MITE amplification, but that this particular MITE copy is probably at the origin of the increase in copy number of the MITE family. On the contrary, the TIR-TE GWAS gave only 51 associations between the presence of a TIR-TE TIP (39 different elements) and a MITE family copy number, but in this case only two of these 39 TIR-TE elements have significant similarity (even limited to the TIRs) with the corresponding MITE family. In these two cases, the region also contains a MITE copy of the same family that could be the genetic factor linked to the family copy number.

All these results suggest that, even if transposases of related class II transposons could mobilize MITEs by a cut-and-paste mechanism, they may not be the main genetic determinant for their amplification. The fact that particular MITE copies show a positive association with MITE copy number may suggests that MITEs amplify by a replicative mechanism of one or few “master” MITE copies, as it is often the case for retrotransposons.

In order to check whether TIP-GWAS reliably identifies “master” copies of replicative TEs, we decided to look for the genetic determinants of rice LINE amplification, as “master” LINEs have particular structural characteristics. Indeed, LINEs are transcribed from an internal promoter located in their 5’ end (Swergold, 1990), absent from the vast majority of elements due to 5’ deletions that make them inactive (Farley et al., 2004). This is the consequence of a transposition mechanism that frequently leads to the integration of elements truncated in 5’, due in part to the low processivity of the RT and to the microhomology-facilitated recombination during integration (Martin *et al*., 2005). Therefore, most newly transposed elements are incapable of expression and transposition, and their structure is different from that of the few “master” elements that are at the origin of each LINE family. It is therefore relatively straightforward to differentiate a “master” LINE from the rest of the copies of the same family.

We performed a GWAS analysis on 104 LINEs families and we found 133 significant peaks corresponding to 79 LINE families (79% of the total families, Supporting Table S6). 57% of the significant peaks (76 peaks from 51 families) had a TIP of the same family at less than 100kb of the corresponding SNP (median distance to SNP = 30.7 Kb, Supporting Table S7). In 99% of these cases, the varieties with the TIP have higher copy number than varieties without the TIP (p < 0.05, two-tailed Wilcoxon rank sum test), suggesting that, indeed, the presence of this particular copy of the TE could be at the origin of the increase in LINE copy number.

We compared the structure of the LINE copies identified as potentially responsible for the amplification of a LINE family with that of the rest of the copies. As an example, Figure 7A (top) shows the Manhattan plot corresponding to the family RIX_comp_MH63_B_R3107_Map6. The three major peaks in chromosomes 8, 11 and 12 identify positions contain a TIP of the same family at positions chr08:5,711,756, chr11:23,993,235 and chr12:1,525,695, respectively. These TIPs (hereafter referred as genetic factors, GF) are in strong linkage disequilibrium (LD) with the minor SNP allele (ie, GF1 and GF2) or with the major SNP allele (ie, GF3), which may explain the association of the SNP with the LINE family copy number. The association of these three LINE insertions with the copy number of this LINE family can also be seen in a TIP-GWAS performed using LINE TIPs as genotype (Figure 7A bottom). Moreover, the presence of the three LINE copies is associated with an increase of copy number of this LINE family (Figure 7B), suggesting that indeed, the three LINE copies may correspond to “master” LINE elements at the origin of this LINE family. In order to validate the presence of these insertions and study the structure of these three potential “master” LINEs, we used the sequence of the Indica varieties from the recently published platinum genomes (Zhou *et al*., 2020). The element corresponding to GF-1 is present in the LIMA::IRGC 81487-1 genome and both GF-1 and GF-3 are present in the genome of the IR 64 variety. A comparison of these elements with all the other copies of the same family present in these genomes shows that they are the only complete ones, the other copies being truncated at 5’ (Figure 7D). All these results suggest that the associations detected point to the active (or “master”) elements that are at the origin of the most recent replicative amplification of these families.

**Figure 7.**
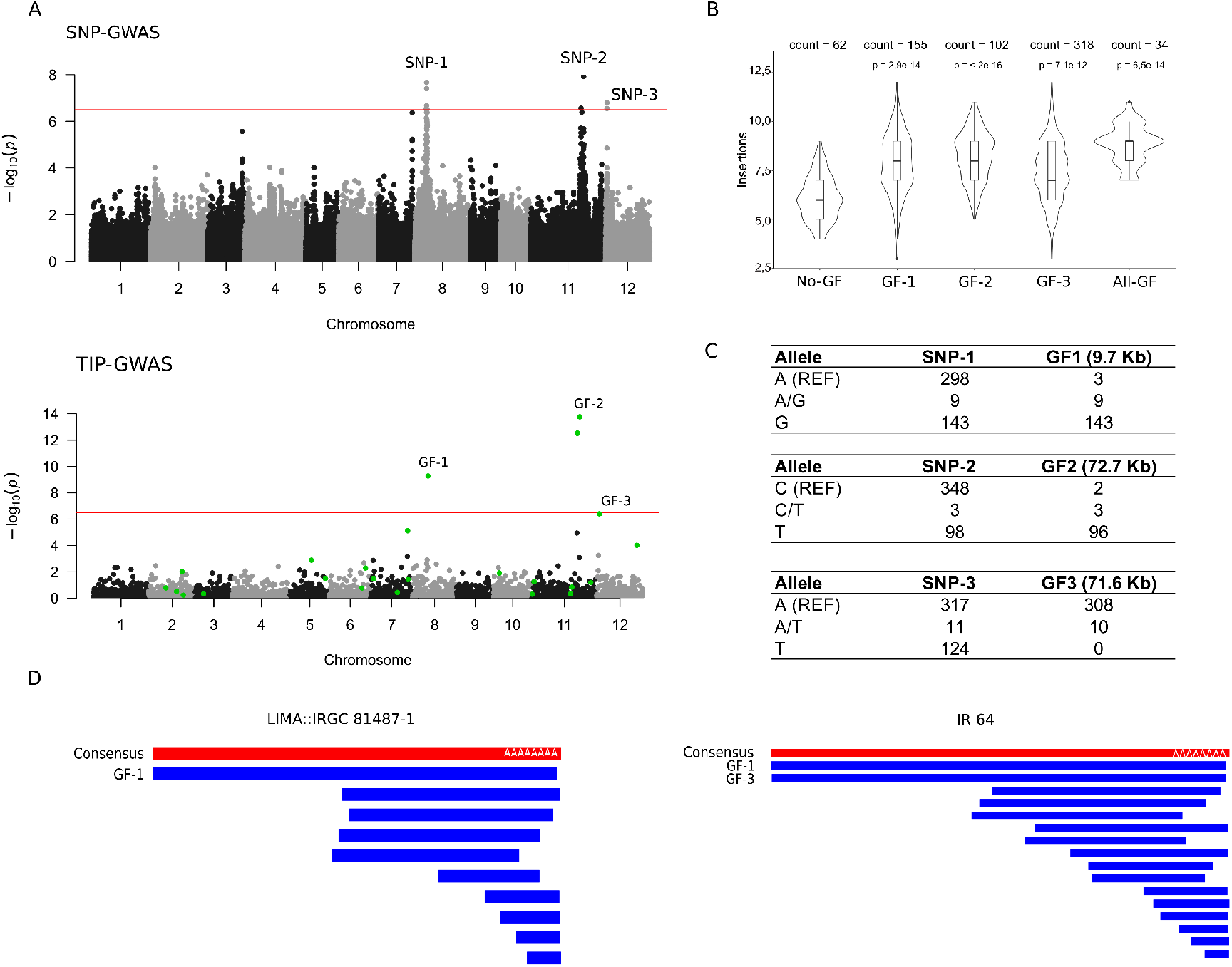
Identification of full-length copies as main genetic factors responsible for LINE amplification in Indica rice. A) SNP-GWAS and TIP-GWAS (451 Indica varieties) analysis of LINE family RIX_comp_MH63_B_R3107_MAP6 using insertion numbers as phenotype. SNPs (upper Manhattan plot) or LINE TIPs (lower Manhattan plot) are used as genotype. Red line represents the Bonferroni-adjusted significant threshold (3.2e^-07^). In green, LINE TIPs of the target family (RIX_comp_MH63_B_R3107_MAP6). GF = Genetic factor) Mean insertion numbers of varieties containing or lacking the putative genetic factors. The number of varieties present in each category are indicated upstream each violin. P-values correspond to two-tailed Wilcoxon rank sum tests comparing each category *vs* the no-GF category. C) Genotypes of the leading SNPs in all Indica varieties and in the subsets of varieties carrying the three different genetic factors. Distance between each GF and the leading SNP is reported in Kb. D) Identification of Genetic Factors in the assembled genomes of LIMA::IRGC 81487-1 and IR64. In red, family consensus sequence. In blue, schematic representation of all genomic BLAST hits obtained using the consensus sequence as query.

These results confirm that GWAS is able to identify the “master” elements at the origin of a replicative amplification of a TE family and strongly suggest that the particular MITE copies found associated with MITE copy number are “master” elements whose replicative amplification is at the origin of MITE families.

### 5- Molecular characteristics of MITEs linked to MITE copy number

As a first attempt to characterize the MITE copies involved in MITE amplification, we compared the sequence of 20 of these MITEs present in the assembled Indica rice LIMA::IRGC 81487-1, as well as of their flanking sequences, with that of the rest of the members of their respective families. We analyzed potential differences in GC content and Minimum Free Energy Structure of these elements including flanks of different length (see methods), as well as TE content of the regions, but failed to detect significant differences between the MITE copies involved in MITE amplification and the rest of the copies of their respective families (Supporting Table S8).

In order to analyze the possible influence of different chromatin characteristics we used the data available for the Nipponbare reference genome to look for the presence in this genome of the MITE TIPs characterized here as associated to the copy number of their respective families. We were able to identify 43 of these MITEs in Nipponbare and we compared them with all the annotated copies of the corresponding MITE families in the whole genome. The characteristics of the chromatin associated to these MITEs seem different than that of the rest of the copies of their respective families. In particular, they show a potential enrichment in transcription activation marks, with 33% more H3K4me3 and 14% more DNAse I hypersensitive peaks coinciding with these specific insertions, as compared with the rest of the copies of their respective families. On the other hand, a recent analysis has characterized the meiotic recombination hotspots in Indica rice mapping them to the Nipponbare genome (Marand *et al*., 2019). An analysis of the overlap of these recombination hotspots with MITE TIPs shows that 18.6 % of the 43 MITEs characterized here as associated to the copy number of their respective families overlap with these recombination hotspots whereas only 12.2 % of all MITEs overlap with these sites.

## Discussion

### MITEs and the evolution of rice genome

The analysis of 103,109 TIPs in 738 rice varieties, which include 382 traditional varieties that represent the variability of the species, allowed us to study the dynamics of the rice mobilome. Our data shows that TEs have been active during the recent evolution of rice and their insertions allow discriminating the different varietal group that have been defined based on SNPs. Interestingly, different types of TEs show a different dynamics in rice. LTR-RTs insertions are present at very low frequency. This result is in agreement with a recent analysis showing that rice LTR-RTs have been active in agro (Carpentier *et al*., 2019), but contrasts with another recent study that found higher population frequencies for LTR-retrotransposons (Kou *et al*., 2020). In line with what the authors of this latter study discuss, we think that these differences are likely due to the inclusion in that study of partial and truncated LTR-RTs and probably old elements (not included in the dataset used here), likely found at higher population frequencies. In contrast to LTR-RTs, MITE show a “U-shape” frequency distribution, with insertions found at low frequency but with an important fraction that seems almost fixed. Almost 40% of the insertions are present in the wild *O. rufipogon* and more than 30% in the relatively distant *O. glumaepatula* genome. This suggests that whereas most LTR-RT insertions are rapidly eliminated, MITE insertions are frequently retained. The smaller size of MITEs, as compared with LTR-RTs, could make their insertions less deleterious and more easily tolerated, and, as MITEs are preferentially found close to genes, the retention of MITEs could simply be due to the difficulty of eliminating them without affecting neighboring genes. However, the fixation of MITEs close to genes could also be the result of a positive selection of some MITE insertions. Our data confirms that MITEs are closely associated to rice genes, concentrating in gene proximal upstream and downstream regions. Many examples of MITE insertions in 5’ and 3’ of genes that alter gene expression in different plants have accumulated over the years, including insertions in promoters enhancing (Zheng *et al*., 2019; Shimada *et al*., 2018; Yin *et al*., 2020; Xu *et al*., 2020) or repressing (Mao *et al*., 2015; Xu *et al*., 2020) transcription, or repressing translation (Shen *et al*., 2017). MITEs have been shown to frequently contain transcription factor binding sites in plants, including rice (Morata *et al*., 2018), suggesting a potential impact on promoter evolution. On the other hand, the methylation of MITEs located upstream or downstream of genes can also repress or activate gene expression, as it has recently been shown for several genes controlling rice tillering (Xu *et al*., 2020). It is therefore possible, that some of the MITE insertions described here may actually modify gene expression, and that some of the insertions present at high frequency in rice may have been positively selected. In particular a number of MITE insertions appear to be present at HF in rice but are absent from *Oryza rufipogon*, the wild species from which rice was domesticated. Therefore, these insertions may have been selected concomitantly with rice domestication. An important fraction of these insertions (66%) are tightly associated to genes and may have altered their coding capacity or their expression. The fact that some of these genes have already been characterized as responsible for important functions linked to rice development and stress responses, suggests that MITEs have played an important role in generating variability used in rice domestication. Future work will be needed to determine the extent of this impact.

Whereas an important fraction of MITEs are fixed in rice, a quarter of the MITE TIPs are present at a population frequency lower than 7% and are probably recent insertions. Up to 61 % of these recent insertions are tightly linked to genes and may therefore be involved in differences of gene expression among varieties and be at the origin of trait variability. In addition, MITEs, as TEs in general, can generate an important number of mutations in relatively short timeframe, which could make their insertions a good alternative to SNPs for GWAS. LTR-RT TIPs have been recently used for GWAS and it was shown that they can uncover additional associations as compared with SNPs (Akakpo *et al*., 2020; Domínguez *et al*., 2020). Here we show that MITE TIPs can also reveal additional associations in GWAS. MITEs are short elements that should be better tolerated than LTR-RTs when inserting close to genes. This may make them a particularly suited source of variability for gene evolution but may also make them particularly suited for markers of gene variability, and therefore, particularly useful for GWAS.

### The mechanism of MITE amplification

MITEs are a particular type of TEs as, although they are deletion derivatives of TIR-TEs which can mobilized them by a conservative cut-and-paste mechanism, they are frequently present in genomes at very high copy numbers. It has been suggested that MITE high copy number may be the result of an increased efficiency of transposition due to their promiscuity in using transposases (Feschotte *et al*., 2003), to particular characteristics of the related transposase in some genomes (Guermonprez *et al*., 2008) or to a higher mobilization efficiency with respect to the related autonomous transposons (Yang *et al*., 2009). Recent genomic studies on the *mPing* MITE have linked its amplification with the presence of a particular copy of the related *Ping* TIR-TE, as this copy is present in varieties showing bursts of *mPing* amplification (Chen *et al*., 2019). On the other hand, an analysis of RILs showing variation in *mPing* copy number, identified QTLs containing multiple copies of *Ping* TIR-TE suggesting a link between *Ping* copy number and *mPing* amplification (Chen *et al*., 2020). Here we have used SNP- and TIP-GWAS to look for genetic determinants of the amplification of 400 MITE families in rice and failed to uncover a link between the presence of particular TIR-TEs and MITE amplification. Although, the mobile nature of TIR-TEs may make it more difficult to reveal this association, these results suggest that, even if transposases of related class II transposons could mobilize MITEs, they may not be the main genetic determinant for their amplification. Interestingly our analyses revealed a positive correlation between the presence in the genome of particular MITE copies and the copy number of the correspondent family, as if only one or few MITE copies were capable of amplifying. This is what commonly happens with replicative TEs, such as LTR-RTs and LINEs, where the replication of one or few “master elements” is at the origin of a whole family. In fact, the phylogenetic analyses of MITE populations in different plant genomes accumulated in the last 20 years (Santiago *et al*., 2002; Naito *et al*., 2009; Xin *et al*., 2019; Lu *et al*., 2012; Feschotte *et al*., 2003) are compatible with the replicative amplification of MITEs, and a replicative transposition mechanism independent of related transposases was proposed long-time ago (Izsvák *et al*., 1999). Izsvak and co-workers proposed that the high secondary structure of MITEs could allow them to fold back to form a stem-loop ssDNA molecule that might detach from the chromosome during replication, in a mechanism similar to that of certain bacterial transposons that move through single-strand transposition associated with DNA replication (Lavatine *et al*., 2016). The fact that only one or few MITE copies are at the origin of a whole MITE family could suggest that the amplification is an efficient but rare event that leads to a substantial increase in copy number at each amplification event. Alternatively, some MITE copies could be more prone to amplification due to their location in the genome or their particular sequence or structural characteristics. The analysis reported here did not reveal major differences between MITE copies positively correlated with an increase of copy number of a MITE family and the rest of the elements of the same family. However, we detected enrichment in chromatin marks associated with active transcription at these elements as well as some enrichment in recombination hotspots compared with the rest of the elements of the family. Origins of replication are closely associated with genes and are enriched in active transcription epigenetic marks (Costas *et al*., 2011), and there are clear links between recombination and replication (Syeda *et al*., 2014). More work will be needed to clarify to what extent these differences may allow some MITE copies to be amplified, but the work presented here is a strong indication for a replicative mechanism of MITE amplification unlinked from the transposase-mediated cut-and-paste mechanism of transposition.

## Conclusions

The results presented here show that an important fraction of MITE insertions is present at high frequency in rice while being absent from its wild ancestor, suggesting that they have been fixed concomitantly with domestication. On the other hand, another fraction of MITE insertions is present at low frequencies among rice varieties, which shows that MITEs have also transposed after domestication. MITEs concentrate close to genes and have generated gene variability during rice domestication and breeding. We used MITE TIPs as genetic information for GWAS and we show that they uncover more associations with rice traits than SNPs. We also used this approach, together with SNP-GWAS to shed light on the still unknown mechanism of MITE amplification. Our results point to the replicative amplification of MITEs by a mechanism uncoupled from their mobilization by class II transposases.

## Methods

### Data source

We used the fastq files from the 1,059 rice varieties sequenced at a minimum coverage of 15X from the 3000 rice genomes project (Li *et al*., 2014), and and three Platinum Indica assemblies (LIMA::IRGC 81487-1, KHAO YAI GUANG::IRGC 65972-1 and LARHA MUGAD::IRGC 52339-1) (Zhou *et al*., 2020)..

### Reconstruction of TE family consensuses

We used TEdenovo from the REPET package (Flutre *et al*., 2011) to build TE consensus sequences from the Japonica Nipponbare genome (Sasaki and Project, 2005), Indica MH63 (Zhang *et al*., 2016) and Aus N22 (Stein *et al*., 2018). TE consensuses were classified at the order level using PASTEC (Hoede *et al*., 2014), and only those classified as “complete”elements were retained. We concatenated the three datasets and removed redundancy by obtaining a library of centroids at 80% identity using VSEARCH (Rognes *et al*., 2016). We run MITE-Hunter (Han and Wessler, 2010) in the three rice genome assemblies and combined the predictions with the MITE dataset from PMITE (Chen *et al*., 2014) and followed the same clustering approach to obtain the MITE library. We concatenated the two libraries (generic TEs and MITEs) to obtain the complete rice TE consensuses (Supporting Dataset S1). Repeatmasker (http://www.repeatmasker.org/) was used to identify genomic regions with similarity to TE consensuses.

### Detection of TE insertions from sequencing reads

We used PopoolationTE2 (Kofler *et al*., 2016) with the mode “separate” to detect TE insertions using the genome of Nipponbare as reference, and we discarded predictions below zygosity of 0.7 as previously recommended (Vendrell-Mir *et al*., 2019). In order to avoid the bias caused by differential sequencing depth, we randomly sampled every accession to 15X prior running the tool using seqtk (https://github.com/lh3/seqtk). To identify non-reference insertions, we intersected all detected insertions with the regions annotated by RepeatMasker and selected all the insertion points that were further than 25bp to any annotated TE in the rice Nipponbare reference genome used here as a reference.

### Estimation of TIP insertion frequencies

Nipponbare genome was split into 500bp (for MITE analyses) and 1Kb (for analyses of the rest of TE) windows. The results of PopoolationTE2 were transformed into bed format. Predicted insertions from different TE orders were separated and intersected with the genome windows to obtain TIPs. TIPs from the 1,059 varieties were concatenated in one file and windows were collapsed to remove redundancy. TIPs of the same order from adjacent windows were merged to obtain a set of high-confidence TIPs. In a second iteration, we intersected all the insertions of the 1,059 varieties, this time excluding the zygosity filter, with the positions of the high-confidence TIP dataset to score the presence/absence status of each TIP in each variety and obtain the final TIP matrices. TIP insertion frequencies were calculated by obtaining the proportion of varieties that contain each insertion.

### Population structure analyses

Principal component analysis was performed using prcomp function from the *stats* R package (The R Foundation for Statistical Computing, 2011). TIP-based Neighbor-joining tree was built using *ape* library from the R package (Paradis *et al*., 2004).

### SNP and TIP-GWAS

We used the LFMM 2 R package (Caye *et al*., 2019) to obtain genotype-phenotype associations applying latent factor mixed models in Indica varieties to correct for population structure (K = 4). For the SNP-GWAS we used the 170 K SNP matrix obtained by Carpentier et al. (Carpentier *et al*., 2019) in lfmm format, derived from the 404k CoreSNP dataset available at SNP-seek (Mansueto *et al*., 2017) (after filtering for minor allele frequency >5% and missing data < 20%). For TIP-GWAS, we used TIP matrices as genotype and considered TIPs with minor allele frequencies > 1%. Bonferroni correction was used to set significance p-value thresholds. Phenotypic traits were downloaded from SNP-seek database.

### Identification of genetic factors

The results of SNP-GWAS using TE family copy number as phenotype and SNPs as genotype were manually inspected to identify peaks with significant associations (Bonferroni-adjusted p value threshold = 3.2e-^07^). We looked for TIPs of the same family located at < 100 Kb, a cutoff based on the patterns of linkage disequilibrium in Indica rice (Mather *et al*., 2007). Statistical significance between family copy number between varieties carrying or not the genetic factor was assessed using two-tailed Wilcoxon rank sum test (p-value cutoff = 0.05).

### Validation of Genetic Factors

Windows containing the genetic factors were extended 4000bp upstream and downstream and extracted from the reference genome (IRGSP, Nipponbare). The regions were mapped to three Platinum Indica assemblies present in our dataset (LIMA::IRGC 81487-1, KHAO YAI GUANG::IRGC 65972-1 and LARHA MUGAD::IRGC 52339-1) using minimap2 (Li, 2018). By screening the best mapping hits we identified the orthologous regions in the three assemblies. Using the family consensus as query for a BLASTN search (Altschul *et al*., 1990) (cutoff e-^20^), we verified the presence of the corresponding TIP in each orthologous region. Schematic representation of all genomic BLAST hits obtained using the consensus sequence as query was performed using Sushi R library (Phanstiel *et al*., 2014).

### Sequence analysis of Genetic factors

The sequence of every MITE in LIMA::IRGC 81487-1 genome was extracted including 100, 250 and 500 bp of upstream and downstream sequences. GC content was determined with bbmap (Bushnell, 2014) and the minimum free energy with ViennaRNA package (Lorenz *et al*., 2011).

### Availability of data and materials

The datasets generated during and/or analyzed during the current study are available in the following repositories: TE consensuses, raw insertions detected by PopoolationTE2 and TIP matrices are available in Zenodo (10.5281/zenodo.4058696); Raw sequencing data of the 1059 rice genomes is available as part of the 3,000 rice genomes project in SRA accession PRJEB6180; H3K4me3 and DNAse I hipersensitivity sites were obtained from GEO database (accession numbers GSM489083 and GSM655033).

## Acknowledgements

RC is recipient of a Juan de la Cierva-formación contract from the Spanish Ministerio de Ciencia y Innovación. This work was supported by grants from the Spanish Ministerio de Economia, Industria y Competitividad (AGL2016-78992-R/FEDER) and Ministerio de Ciencia y Innovación (PID2019-106374RB-I00 /AEI / 10.13039/501100011033) to JMC. Computing resources have been provided by the Red Española de Supercomputación at the Pirineus machine (RES activity BCV-2019-2-0006). We thank Marie Mirouze (Université de Perpignan) for helpful discussions.

## Authors’ contributions

JMC and RC designed the analysis. RC, PV-M and AB performed TIP detection and analyzed the data. RC performed GWAS analyses with the collaboration M-C C and OP. JMC and RC wrote the manuscript with collaborations of PV-M and AB. All authors revised and approved the manuscript.

## Legends for Supporting Information

**Supporting Figure S1**. Percentage of the TE library occupied by each TE order. DHX = Helitrons, DTX = TIR-TE transposons, RIX = LINEs, RLX = LTR-retrotransposons, RSX = SINEs.

**Supporting Figure S2**. Insertion detection bias between CAAS and IRIS varieties. Number of PoopulationTE2 filtered insertions per variety in 1059 coverage-homogenized genomes.

**Supporting Figure S3**. Neighbor-joining trees based on TIPs from six different TE groups. DHX = Helitrons, DTX = TIR transposons, RIX = LINEs, RLX = LTR-retrotransposons, RSX = SINEs.

**Supporting Figure S4**. Genome wide density plot of MITE and LTR-retrotransposon TIPs.

**Supporting Figure S5**. Genome wide density plot genes, MITEs and LTR-retrotransposons according to their High (Q1) or Low (Q4) frequency on the population.

**Supporting Figure S6**. A) Non-reference insertions per 100Kb of feature (738 varieties). Each boxplot represents the distribution of mean insertions per variety in each feature normalized by the total length of feature in the whole genome. B) Non-reference insertions per 100Kb of feature considering by high- and low-frequency insertions (Q1 and Q4 quantiles, respectively).

**Supporting Figure S7**. Grain width of varieties carrying MITE-1 insertion (chr05:5364000-5365000) and the deletion located 4,500 bp upstream the GSE5 gene described in (Duan et al. 2017).

**Supporting Dataset S1**. Consensus sequences of the 821 TE families.

**Supporting Table S1**. Description of the 1,059 varieties used in this study.

**Supporting Table S2**. List of High-frequency MITE insertions present in *O sativa* ssp. *Japonica* cv. Nipponbare and absent in *O. rufipogon*, present in genic regions.

**Supporting Table S3**. List of SNPs and TIPs significantly associated with grain width.

**Supporting Table S4**. List of MITE TIPs significantly associated with five rice phenotypes.

**Supporting Table S5**. List of 18 MITE families and their linked transposases showing significant similarity on the 5’ and 3’ extremities, along with the results from the SNP-GWAS and TIP-GWAS using TIR-TEs and MITEs.

**Supporting Table S6**. GWAS leading SNPs associated with family copy number for 428 TE families.

**Supporting Table S7**. Information related to the GWAS leading SNPs and TIPs associated with family copy number.

**Supporting Table S8**. GC content, Minimum Free Energy Structure and TE content (250 Kb region) of GF and non-GF loci of 25 MITE families present in LIMA::IRGC 81487-1 Genome.

