## Supplemental Figure S1 for "The replicative amplification of MITEs and their impact on rice trait variability"

### Slide 1
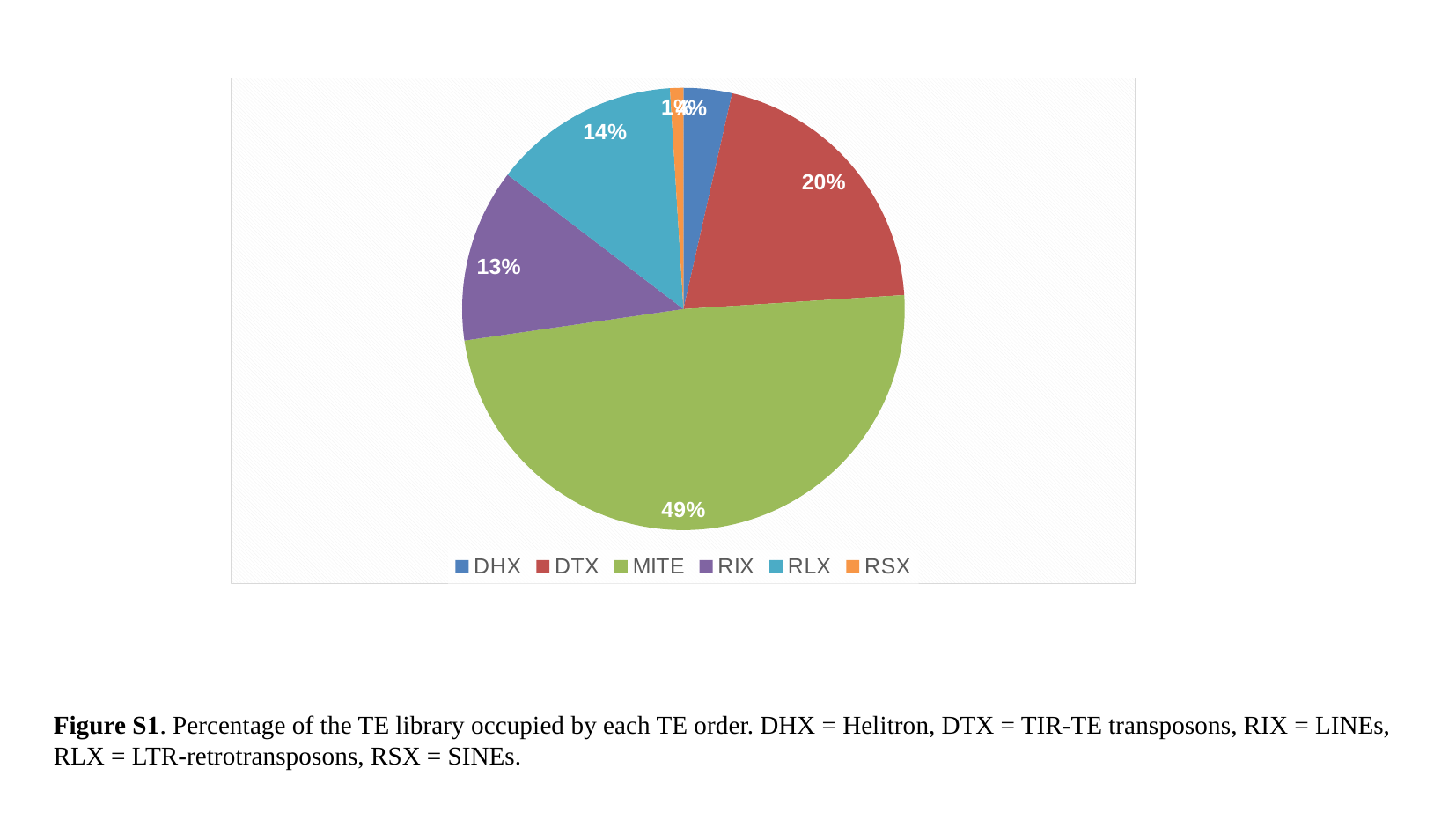

#### Chart
| Category | Custom library |
|---|---|
| DHX | 29.0 |
| DTX | 168.0 |
| MITE | 400.0 |
| RIX | 104.0 |
| RLX | 112.0 |
| RSX | 8.0 |Figure S1. Percentage of the TE library occupied by each TE order. DHX = Helitron, DTX = TIR-TE transposons, RIX = LINEs,
RLX = LTR-retrotransposons, RSX = SINEs.
