## Supplemental File S2 for "The replicative amplification of MITEs and their impact on rice trait variability"

### Slide 1
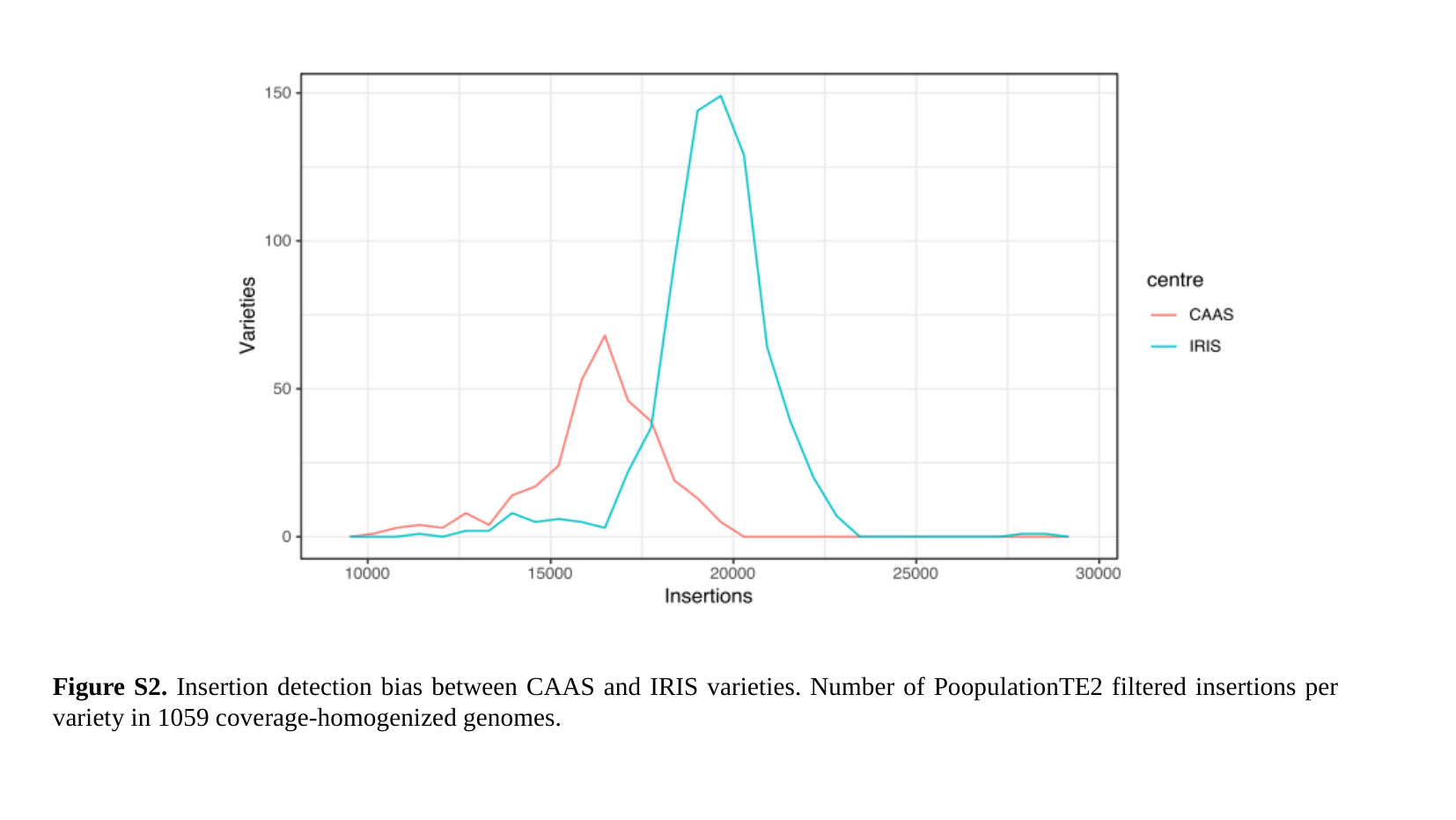

Figure S2. Insertion detection bias between CAAS and IRIS varieties. Number of PoopulationTE2 filtered insertions per variety in 1059 coverage-homogenized genomes.
