## Supplemental Figure S3 for "The replicative amplification of MITEs and their impact on rice trait variability"

### Slide 1
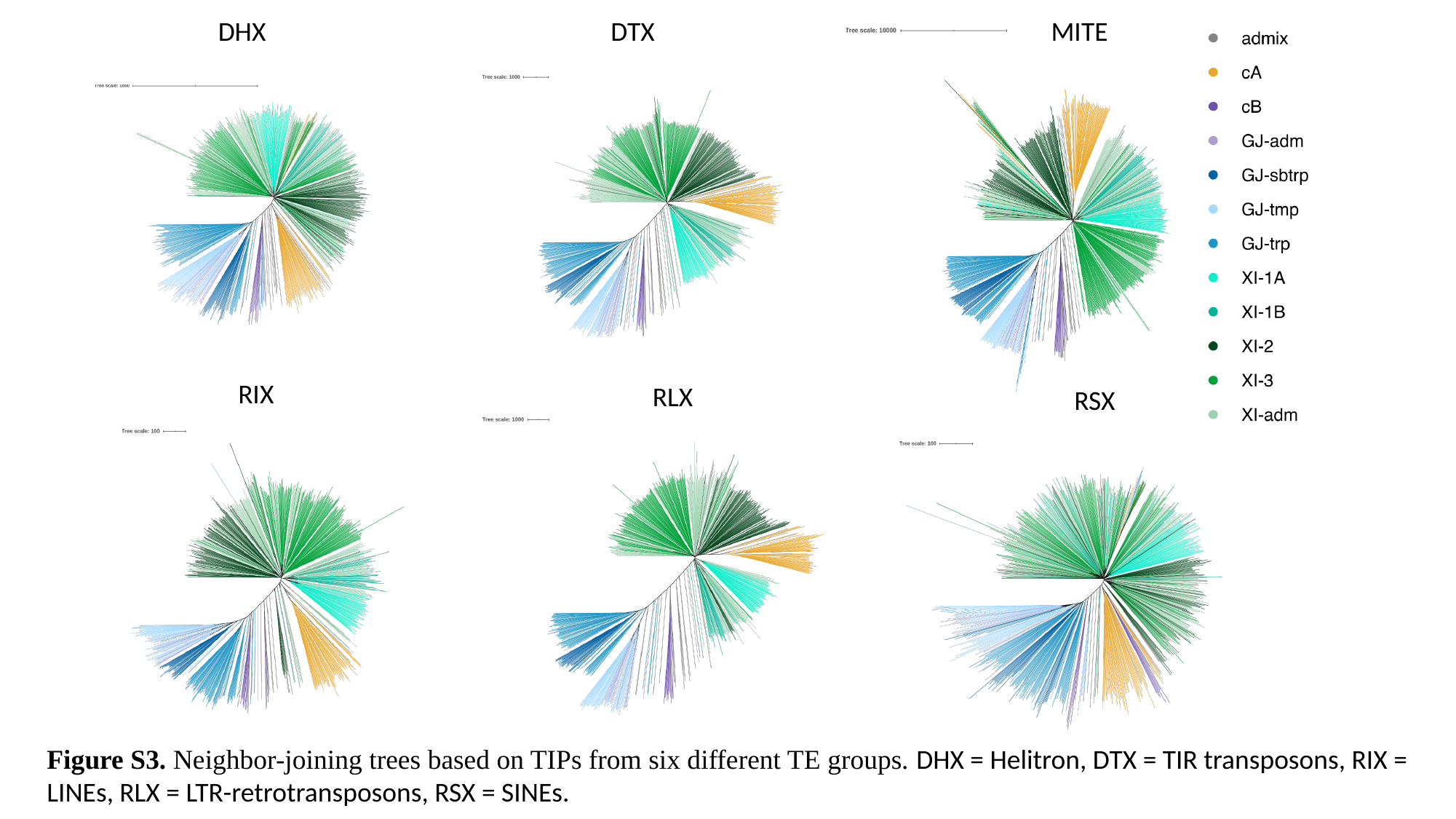

DHX
DTX
MITE
RIX
RLX
RSX
Figure S3. Neighbor-joining trees based on TIPs from six different TE groups. DHX = Helitron, DTX = TIR transposons, RIX = LINEs, RLX = LTR-retrotransposons, RSX = SINEs.
