## Supplementary figures and images for "The replicative amplification of MITEs and their impact on rice trait variability"

### Supplemental Figure S4

#### Slide 1
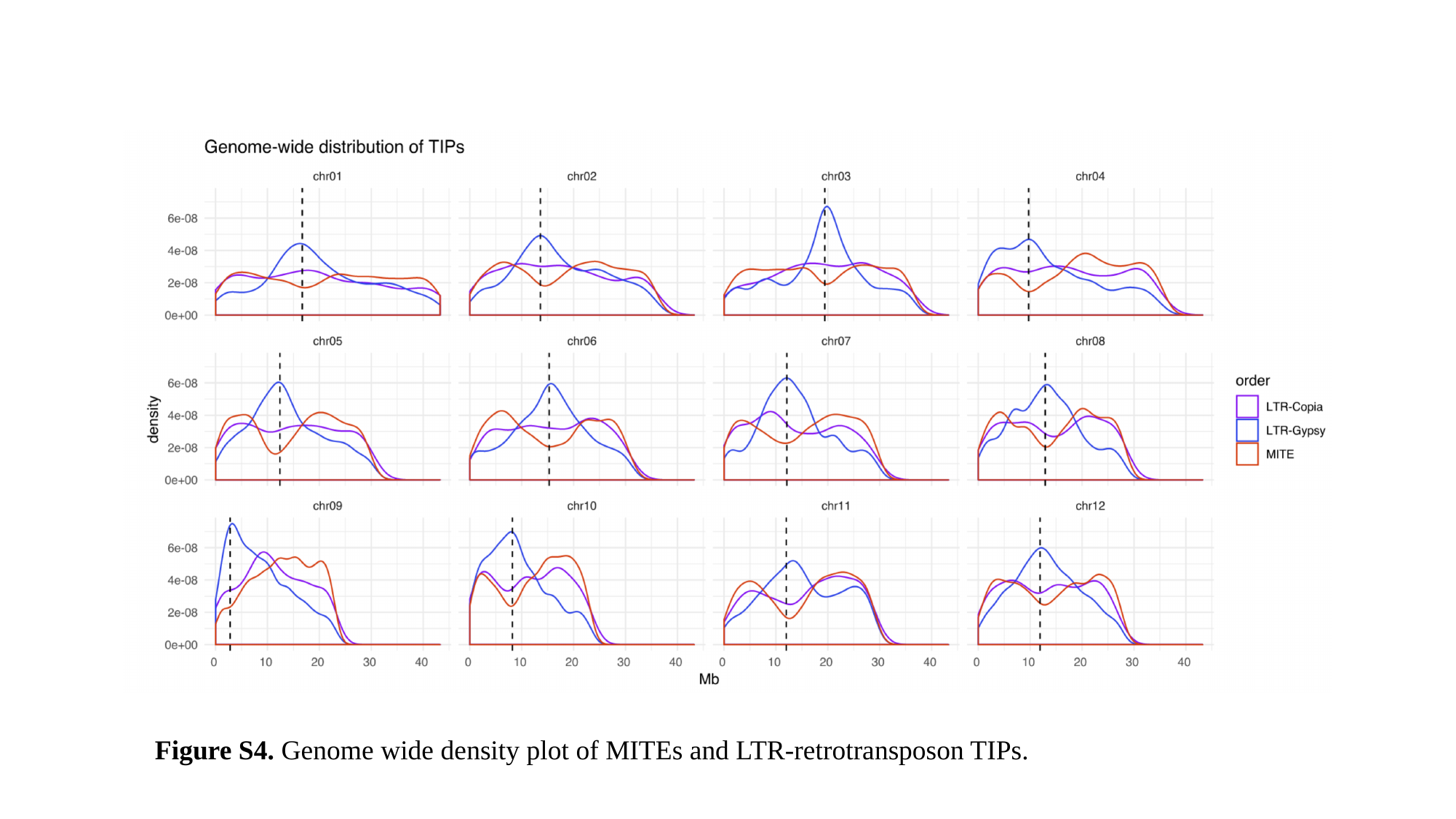

Figure S4. Genome wide density plot of MITEs and LTR-retrotransposon TIPs.
