## Supplemental File S5 for "The replicative amplification of MITEs and their impact on rice trait variability"

### Slide 1
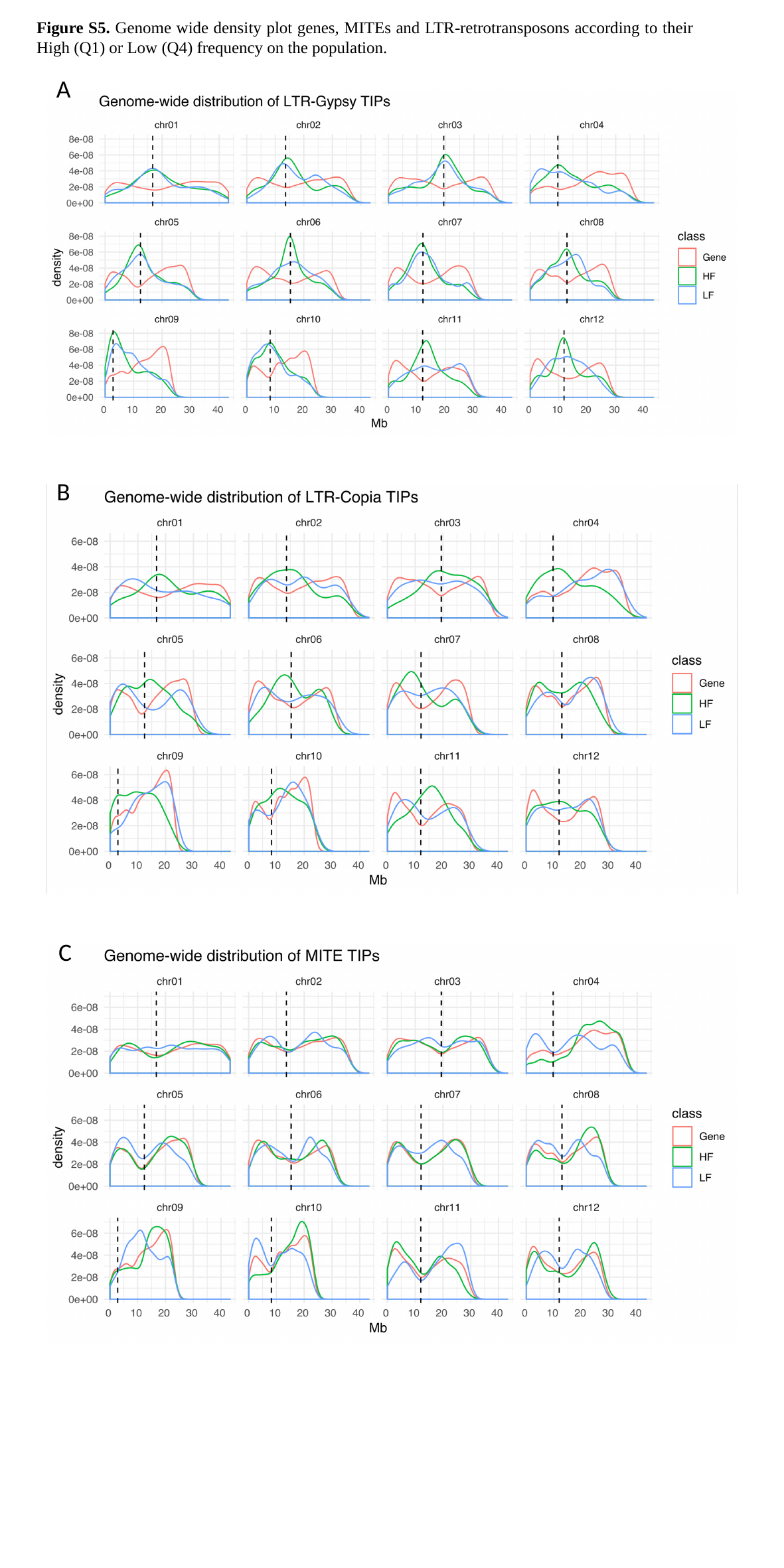

Figure S5. Genome wide density plot genes, MITEs and LTR-retrotransposons according to their High (Q1) or Low (Q4) frequency on the population.
A
B
C
