## Supplemental File S6 for "The replicative amplification of MITEs and their impact on rice trait variability"

### Slide 1
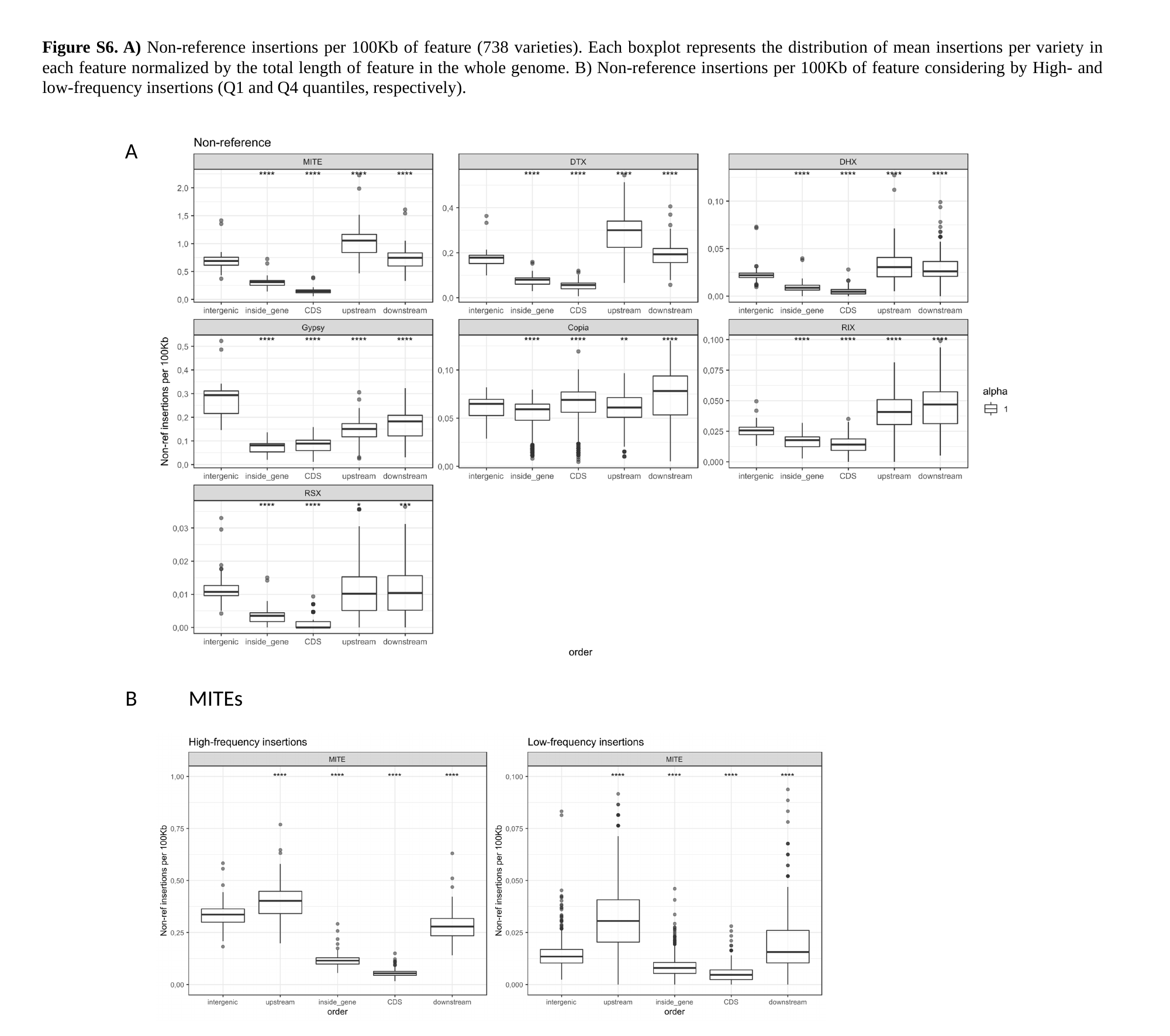

Figure S6. A) Non-reference insertions per 100Kb of feature (738 varieties). Each boxplot represents the distribution of mean insertions per variety in each feature normalized by the total length of feature in the whole genome. B) Non-reference insertions per 100Kb of feature considering by High- and low-frequency insertions (Q1 and Q4 quantiles, respectively).
A
B
MITEs
