## Supplemental File S7 for "The replicative amplification of MITEs and their impact on rice trait variability"

### Slide 1
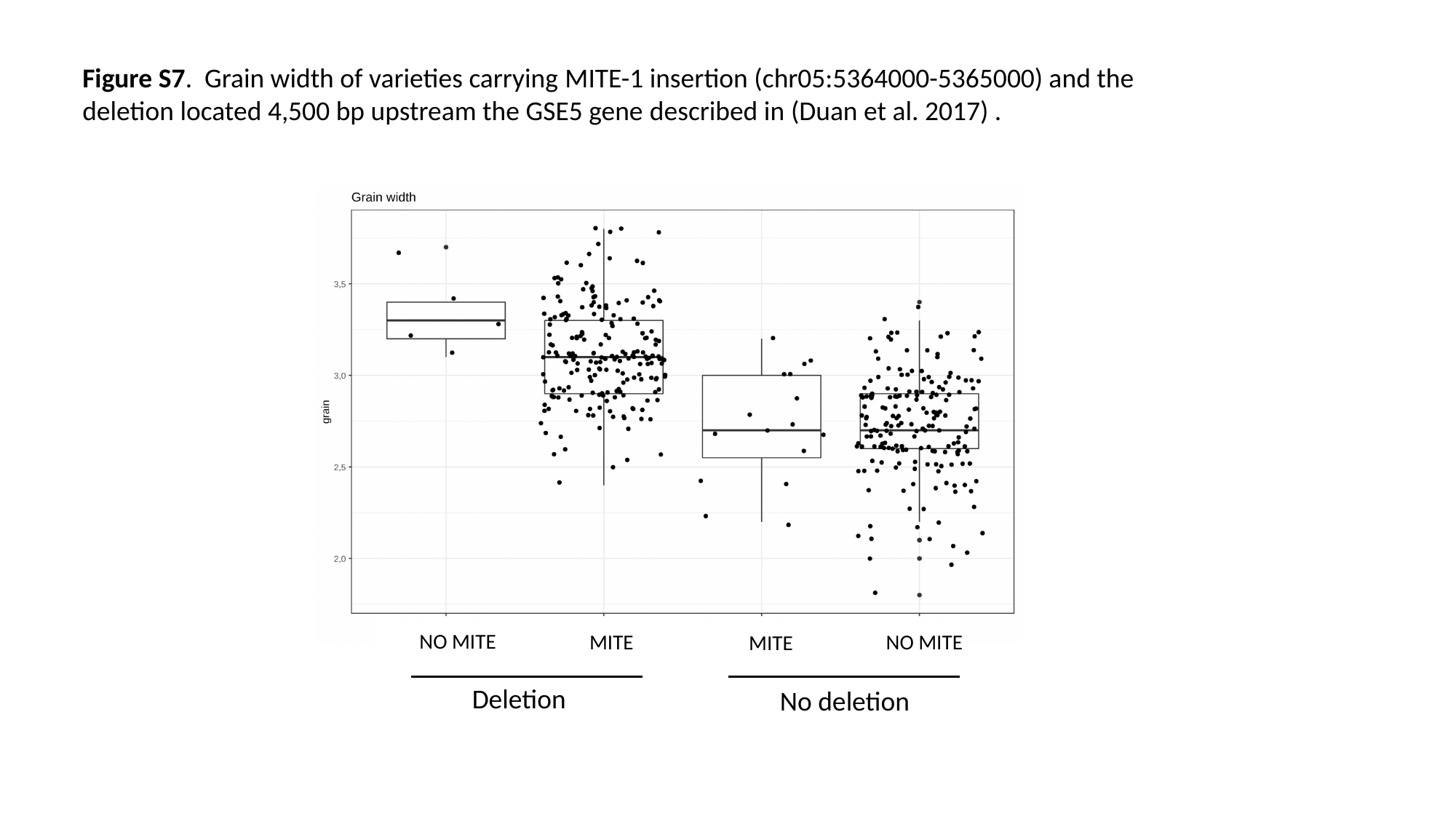

Figure S7. Grain width of varieties carrying MITE-1 insertion (chr05:5364000-5365000) and the
deletion located 4,500 bp upstream the GSE5 gene described in (Duan et al. 2017) .
NO MITE
MITE
NO MITE
MITE
Deletion
No deletion
